# Modeling the Cell Cycle Response to Carbon and Nitrogen deprivation in *Caulobacter* Populations

**DOI:** 10.1101/2022.07.23.501216

**Authors:** Chunrui Xu, Bronson R. Weston, Yang Cao

## Abstract

*Caulobacter crescentus* inhabits a wide range of aquatic ecosystems, including environments with poor nutrients. It undergoes an asymmetrical cell division cycle, generating a pair of daughter cells with distinct motility and replicative potentials. *Caulobacter* populations have the flexibility to save energy by halting chromosome replication and reduce intraspecific competition by settling in different places in environments. The control mechanisms underlying *Caulobacter* cell development have been well studied under nutrient-rich conditions, however, its mechanism of response to stressful changes is not fully understood. Here we present a mathematical model to analyze the starvation responses in *Caulobacter*. We investigate several known starvation signaling pathways to study how these pathways influence cell cycle development and explain experimental observations of starved *Caulobacter* populations. We also apply a new parameterization strategy to mathematical modeling of biological systems, whose diverse communities have to be robust with many parameter variations, while still having accurate control to maintain regular cell cycle dynamics. Our model demonstrates that the guanine-based second messenger, c-di-GMP (cdG), plays important roles to immediately arrest the cell cycle of *Caulobacter* under nutrient deprivation; however, it is not sufficient to cause the robust arrest. Our model suggests there should be unknown pathway(s) reducing the levels of CtrA under starvation condition, which results in a significant delay in cytokinesis of starved stalked *Caulobacter* cells.

## 1 Introduction

*Caulobacter crescentus* utilizes a dimorphic lifestyle to survive in nutrient-poor environments [16, 1, 13]. It divides asymmetrically, producing two different daughter cells [10] (Fig. 1 A). One has a stalk to attach to solid surfaces in environments, called ‘stalked’ cell, which is non-motile and ready to replicate DNA. The other (called ‘swarmer’ cell) develops pili and a flagellum to move away and search for a satisfactory place in environments. The swarmer cell can not replicate DNA until it differentiates into a stalked cell morphology given favorable environments [7].

**Figur 1:**
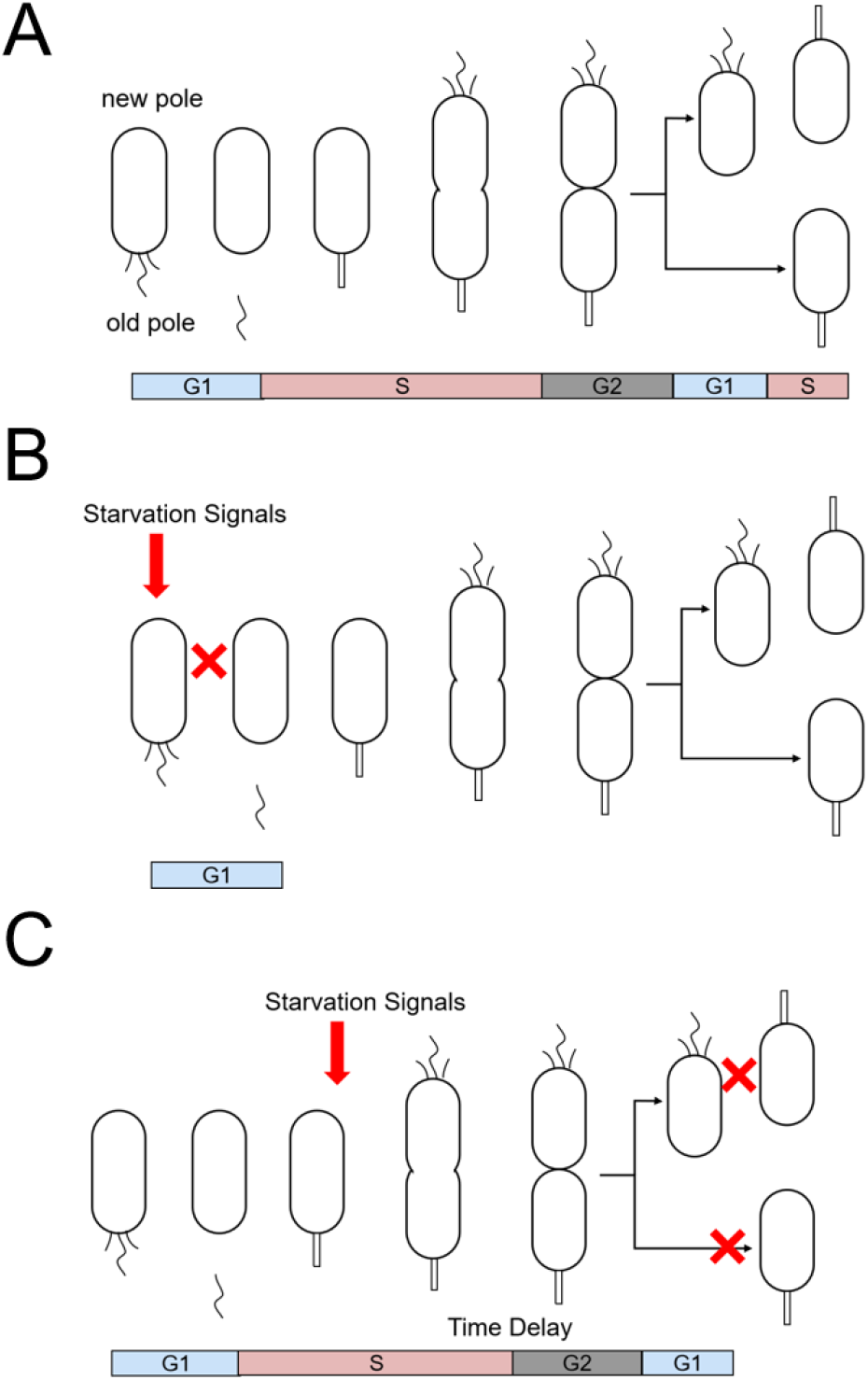
Nutrients starvation leads to G1 arrest. (A) The normal asymmetrical cell cycle of *C. crescentus* in nutrient-rich conditions. (B) Starved swarmer cells show immediate G1 arrest. (C) Starved stalked cells show delays in S phase and arrest at the second G1 phase.

The asymmetrical division cycle of *Caulobacter* requires complex coordination of genetic expression, proteolysis, phosphorylation, and second messenger signaling. System biologists have proposed a series of mathematical models to study the temporal regulations of chromosome replication and cell differentiation of *Caulobacter* [14, 15, 19, 20, 22]. Li et al. [14, 15] and Weston et al. [21] have utilized a core cell cycle regulatory network, consisting of four master regulators CtrA, DnaA, GcrA, and CcrM, to investigate the mechanisms and capture the behaviors of the cell cycle development in wild type and novel mutant cells. For cells living in natural environments, nutrition changes are inevitable and have to be handled efficiently. Xu et al. have preliminarily studied how second messengers detect and respond to the environmental nutrition changes through PTS systems [22]. However, how the *Caulobacter* cell cycle responds to starvation and the corresponding detailed mechanisms are yet elucidated. In this study, we aim to utilize mathematical modeling to study the control mechanism of cell cycle in the context of nutrient stress.

Under nutrient starvation, *Caulobacter* swarmer cells show rapid and robust G1 arrest, conserving energy by halting DNA replication and keeping motility to search for nutrients [6, 3, 17] (Fig. 1 B). Starved stalked cells go through significant delays in cell division, and eventually arrest in G1 phase (Fig. 1 C). To explain the control mechanisms underlying starved cell development, we commence building the model from integrating our previous second messenger model [22] into a temporal cell control model [21], where the vital regulatory second messenger cdG plays important roles to receive and respond to environmental cues and influences the activity of downstream regulator CtrA through multiple pathways. We further include other known cdG- and CtrA-relevant pathways, which are involved in stressful regulations into the model, such as the cdG-ShkA-TacA-SpmX pathway [9], RpoD-regulated transcriptions [11, 4], and hierarchical assembly of ClpXP complexes [8]. Additionally, the chromosome replication initiator protein DnaA is reported to be dramatically reduced under starvation, which is likely dependent on antagonists of cdG - second messengers (p)ppGpp [12]. Experiments also suggest that the cell growth of *Caulobacter* would slow down under starvation stress [6, 17].

## 2 Methods

In this study, we integrate our previous second messenger model [22] into Weston et al.’s temporal model [21] with the connections between second messengers and the master regulator CtrA. Additionally, we modified and improved Weston et al.’s model with newly investigated pathways to include known stressful response pathways and enhance the biological accuracy (Fig. 2). Here we specify our major modifications.

**Figur 2:**
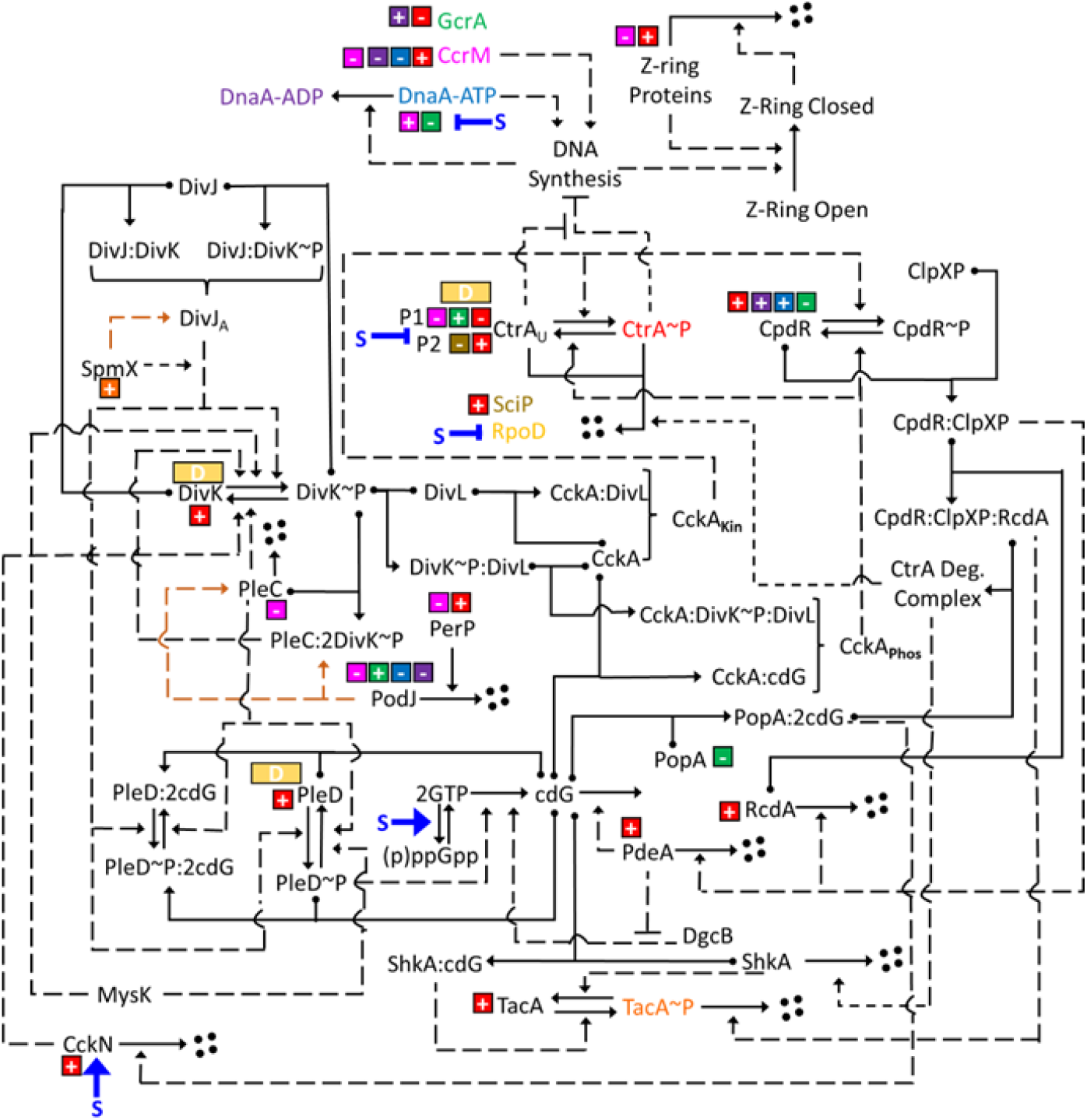
Diagram of regulatory interactions and starvation signals captured by this study. A box with a plus or minus sign indicates whether the protein indicated by the same color activates or inhibits expression of the gene. The sigma factor RpoD is indicated as yellow where its promoter regulation is indicated in a yellow box. Dot-headed solid line indicates the binding of species. Chemical conversion of one species to another is indicated by an arrow. An arrow from a protein to four black circles represents Proteolysis. Dashed arrows indicate an ‘influence’ of a protein on a chemical reaction. A bright blue S indicates a starvation signaling, with arrow for activation and bar for inhibition).

### Modeling PTS^Ntr^/SpoT nutrient signaling cascade through cdG

Xu et al. [22] modeled the response of second messengers - (p)ppGpp, GTP, and cdG - to starvation signals in *Caulobacter*. The nitrogen phosphotransferase system (PTS^Ntr^) detects external nutrient levels and transfers signals to downstream bifunctional enzyme SpoT. Under starvation, SpoT dramatically influences GTP levels by regulating the conversion between GTP/GDP and (p)ppGpp, and leads to reduced cdG levels in sequence. Here, we model the fluctuation of GTP levels under different conditions by combining Xu et al.’s second messenger network [22] with Weston et al.’s cell cycle control model to add a coefficient 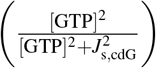 on the synthesis simulation of cdG. *J*_s,cdG_ = 1500*µM* [22] indicates the concentration of GTP corresponding to half of maximal synthesis. Under nutrient-rich conditions, [GTP] *≈* 1221*µM*; under nutrient-starved condition, [GTP] *≈* 227*µM* [22]. Therefore, the fractional term 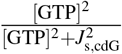 is reduced to only 5.6% under starvation. Thereby, we can analyze the impact of environmental nutrition changes through cdG-dependent pathways on *Caulobacter* cell development.

### cdG regulates the morphogenesis of *Caulobacter* through the ShkA-TacA-SpmX pathway

cdG binds to ShkA to stimulate its kinase activity [2, 5]. Phosphates are transferred from ShkA kinase to the phosphotransferase ShpA, and then received by TacA in sequence [2, 9]. Phosphorylated TacA controls the transcription of SpmX, while the latter recruits and activates DivJ to the stalked pole (old pole, Fig. 1 A). As DivJ is a key kinase regulating the phosphorylation state of regulators such as DivK and CtrA, this section connects starvation signaling with phosphotransfer in *Caulobacter* cells (see Figure. 2). The modeling of ShkA-TacA-SpmX-DivJ pathways is shown in Supplementary File 1.

### Modeling the hierarchical assembly of ClpXP complexes

ClpXP is a conserved protease degrading a wide range of proteins in *Caulobacter*, such as CtrA and ShkA. Although ClpXP levels are almost constant over the cell cycle, it hierarchically assembles helpers to cyclically degrade proteins [18]. Here, we model the hierarchical assembly of ClpXP complexes, [ClpXP:CpdR], [ClpXP:CpdR:RcdA], and [ClpXP:CpdR:RcdA:PopA:cdG2] (named as ClpXP complex in the Supplementary File 1), in order, where [ClpXP:CpdR:RcdA] is not modelled in Weston et al.’s model [21].

### Modeling RpoD-regulated transcriptions

The sigma factor RpoD (*σ*^70^) in *Caulobacter* directly mediates the transcription of housekeeping genes, while sigma factors are observed to respond to stresses and influence downstream gene transcriptions in other bacteria, such as *E. coli* and *B. subtilis* [4]. RpoD levels reduce to 78% in 30 min after carbon starvation in *Caulobacter*, suggesting a RpoD-dependent pathway of stringent response [4]. Based on the observations of RpoD-targeted proteins that DivK levels, CtrA levels, and PleD levels decrease to 70%, 40%, and 10% in wild type cells upon 60 min carbon deprivation, respectively, we introduce ‘RpoD’ as an adjustable parameter, being 1 in nutrient-rich condition and 0.1 in starved condition, into the modeling of synthesis of DivK, CtrA, and PleD (See Supplementary File 1).

### Parameterization algorithm

Here we utilize a parameterization algorithm, the Monte-Carlo Markov-Chain, which was first used in Weston et al. [21]. In our study, a list of mutant strains is used to estimate parameters: P*pleC*::Tn, Δ*ccrM*, Δ*gcrA*, Δ*pleD, divL*A601L, *ctrA*Δ3Ω, *ctrA*D51E, P*pleC*::Tn&Δ*divJ*, Δ*divJ*.

The main reason for this algorithm is that biological cells, even those of the same type, demonstrate great varieties. The traditional öne set of parameters fits allstrategy is not realistic. For parameter estimation, we first choose a ‘current’ parameter set *p* = (*p*_1_, *p*_2_, …, *p*_*m*_) of length *m* and perturb it to generate a ‘temporary’ parameter set *p*^*′*^, such that:

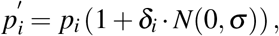

where *N*(0, *σ*) is a random number drawn from a normal distribution with mean zero and standard deviation *σ*. *δ*_*i*_ is a Boolean variable, that denotes whether or not a given parameter *p*_*i*_ will be subjected to random perturbation. Additionally, any 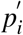 that evaluates to a negative number is set to 0.

In every iteration, we define the probability that a temporary parameter set replaces the ‘current’ parameter set as

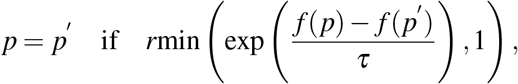

where *r* is a random number selected from a uniform distribution between 0 and 1, *f* is the cost function for the parameter set, and *τ* is the ‘cooling factor’ for simulated annealing. Thus, any temporary parameter set that has a lower cost than the ‘current’ parameter set replaces the ‘current’ parameter set. Any temporary parameter set that has a higher cost than the ‘current’ parameter set has a probability *Pr* of replacing the current parameter set. This process of generating a new (temporary) parameter set and comparing it with the current parameter set is considered one iteration (one step) of the random walk in the parameter space.

The efficiency of the algorithm depends on the parameters *σ, τ* and *n*. A smaller *σ* leads to smaller perturbations. If *σ* is small, it is likely to generate a *p*^*′*^ that is an improvement over *p*. However, it may take much longer to make large improvements. *τ* dictates the probability that a *p*^*′*^ that has a higher cost than *p* will be accepted as the new p. A larger *τ* improves the algorithm’s chances of escaping a local minimum but decreases the probability of identifying a new local minimum. When n is large, more parameters are perturbed in each iteration. As a result, each *p*^*′*^ will be further away (in parameter space) from *p*. The advantage to a large *n* is that fewer steps are needed to make a large change; however, the chance that *p*^*′*^ is an improvement over p decreases.

### Cost function

The cost function *f* (*p*) is the sum of the scores of simulations for a wild-type cell and selected mutant strains:

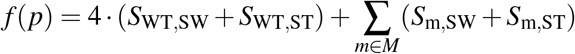

where *M* is the set of mutant strains used for parameterization.

The score of WT cells is defined in Weston et al’s model [21]:

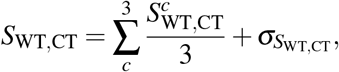

where CT (cell type) is either SW or ST cell type. *c* denotes the index of cycle (the last three cycles in 1600 min simulation are counted). *σ* indicates the standard deviation between individual scores. 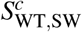 consists cost scores for gaps of concentrations, replication initiation time, and division time between simulations and experimental observations, while 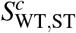 consists cost scores for gaps of replication initiation time, and division time between simulations and experimental observations (shown below).

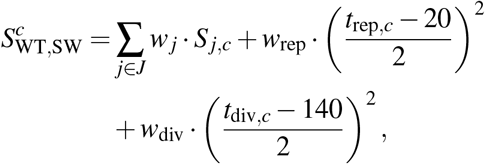

where *J* is a collection of experimental data and *j* is the index of one set of experimental data. *w*_*j*_, *w*_rep_, and *w*_div_ are weights of concentration gap, DNA replication time gap, and division time gap between simulations and experiments, respectively.

*S*_*j,c*_ measures the agreement between the simulation and experiment:

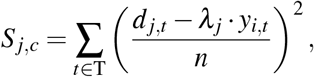

where *T* is a collection of time points and *t* is one given time point. *d*_*j,t*_ indicates the normalized experimental concentration of the *j* experiment at time *t*, while *y*_*j,t*_ is the corresponding simulated value. *n* is the number of data points and *λ*_*j,t*_ is a scaling coefficient.

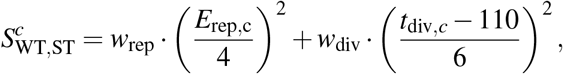

where *E*_rep,c_ is the error associated with estimated chromosome replication initiation time of ST cells.

Additionally, we incorporate mutant DivK*∼*P data into our cost function to guide our model into accurate parameter sets for DivK*∼*P, which is not considered in Weston et al’s model [21]. The scores of P*pleC*::Tn and Δ*divJ* mutant strains are modified in this study as follows:

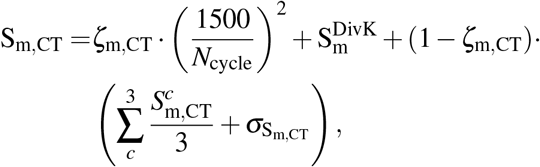

where *ζ*_m,CT_ is the arrest coefficient for strain m and equals to one if the cell cycle is arrested and zero if not. 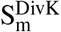 is the DivK related score for strain m, 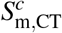 is the cell cycle specific score.

Cell cycle specific scores of P*pleC*::Tn and Δ*divJ* are same with Weston et al [21]:

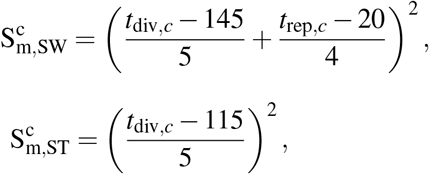

where *t*_div,*c*_ and *t*_rep,*c*_ indicate the time of division and chromosome replication in cell cycle *c*.

DivK specific scores are calculated as:

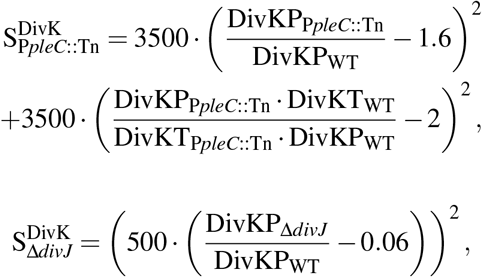

where DivKP_m_ and DivKT_m_ are the average DivK*∼*P levels and total DivK levels, respectively, for strain m.

The cost functions of other mutant strains (Δ*ccrM*, Δ*gcrA*, Δ*pleD, divL*A601L, *ctrA*Δ3Ω, *ctrA*D51E, and P*pleC*::Tn&Δ*divJ*) are provided in Supplementary File 2.

## 3 Results

### 3.1 Our model matches well with wild type and mutant strains in nutrient-rich condition

To ensure our new model and parameter sets are in agreement with experimental observations, we analyze the prediction accuracy of 35 mutant strains viability (both for swarmer and stalked cells) with 150 parameter sets. The success rate of prediction is about 80%, which is reasonable considering the complexity of this model and a large number of predicted cell types. Additionally, we compare our simulations of a wild type (WT) cell with experimental data (Fig. 3). The simulated temporal dynamics of key regulators in a WT swarmer cell fit quite well with experiments.

**Figur 3:**
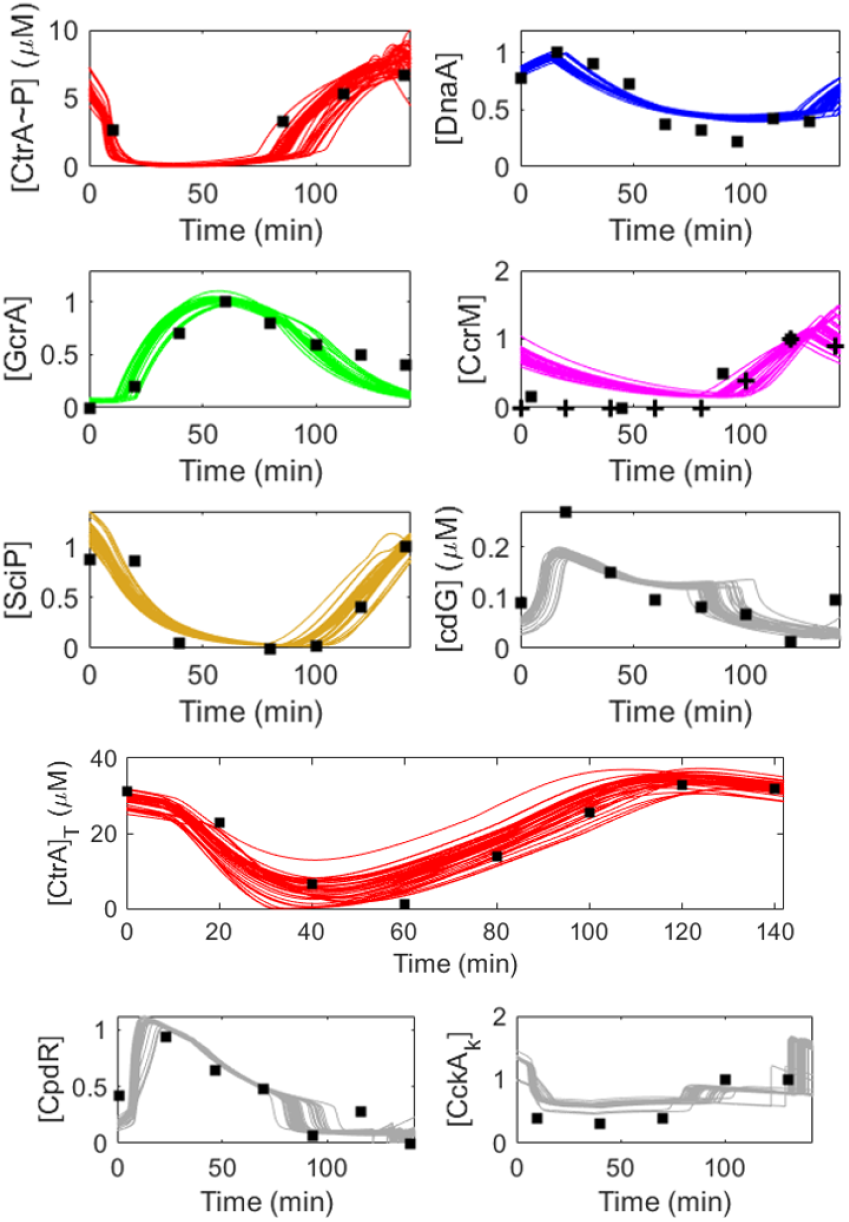
Model and parameter sets fit experimental data reasonably well. For 50 parameter sets that are randomly chosen from 200 parameter sets obtained by optimization, we plot swarmer-cell simulations in comparison to experimental data. Data are collected as follows: CtrA*∼*P: Jacobs et al. 2003; DnaA: Cheng and Keiler 2009; GcrA: Holtzendorff et al. 2004; CcrM: plus from Zhou and Shapiro 2018; square from Grunenfelder et al. 2001; total CtrA: Mcgrath et al. 2006; CpdR: Iniesta et al. 2006; CckA kinase: Jacobs et al. 2003.

### 3.2 Introducing known starvation signals (Signal 1) successfully captures the G1 arrest of *Caulobacter* population, but the delayed cytokinesis of starved stalked cells requires reduced CtrA levels

Given reasonable agreement of our model with experiments, we then apply this model to investigate how our model responds to environmental signals. We first introduce a starvation signal defined as ‘Signal 1’ (Table 1) and compare simulations with observations (Fig. 4). Our simulation with Signal 1 indicates that the introduced starvation pathways are sufficient to arrest swarmer and stalked cells at the G1 stage, while the observed delay in stalked cell division under starvation is not captured (Fig. 4 A, B). Additionally, simulation with signal 1 fails to capture reduced levels of CtrA under starvation, which suggests that there might be an unknown starvation signaling pathway decreasing CtrA levels (Fig. 4 C). We further introduce Signal 2 (Table 1), which enforces the reduction of CtrA expression on Signal 1. Our results then show good agreements with experiments (Fig. 5), indicating that the reduced CtrA expression is required to delay the stalked cell division when nutrients are deprived.

**Tabell 1:**
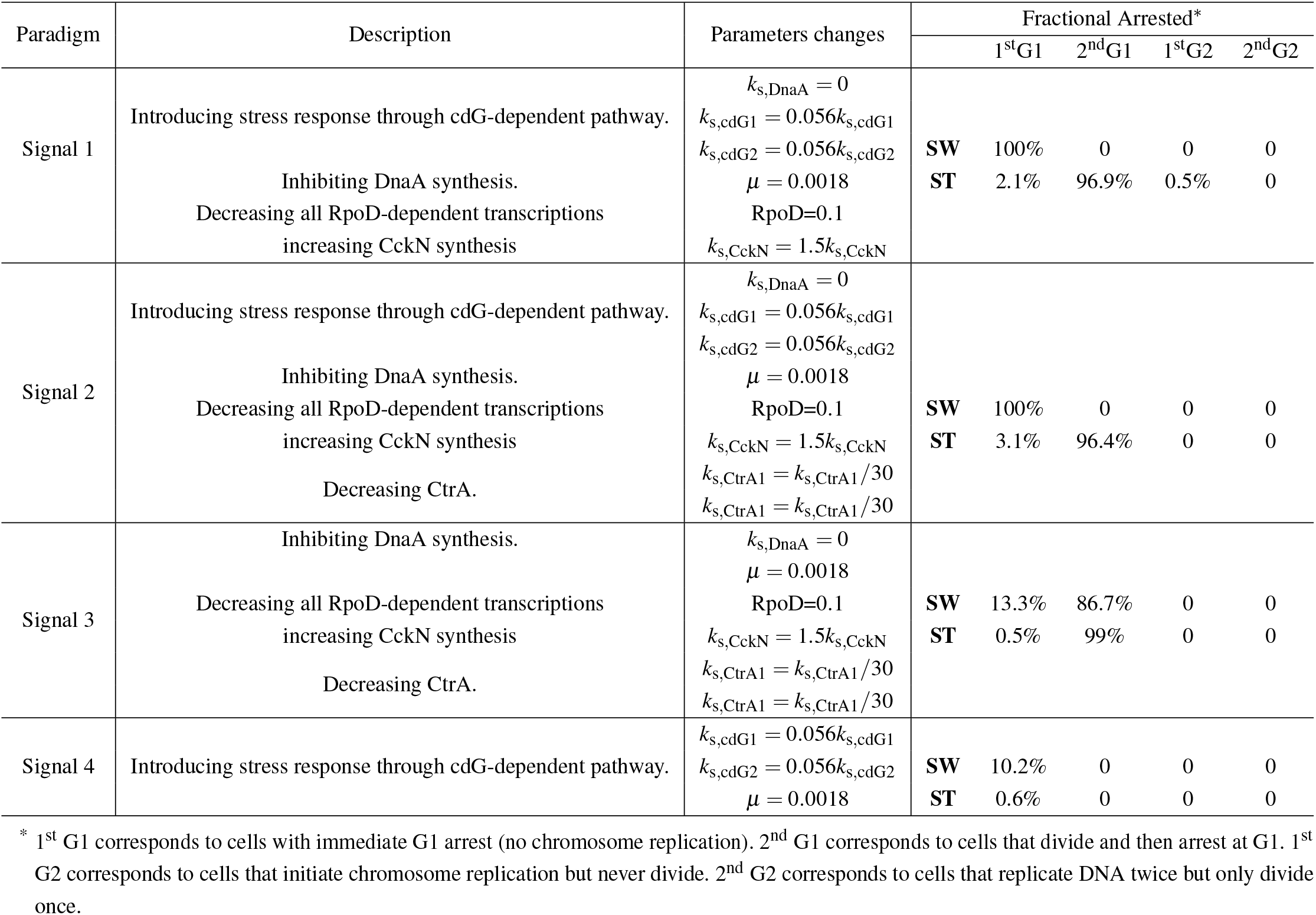
Signaling targets and arrest statistics.

**Figur 4:**
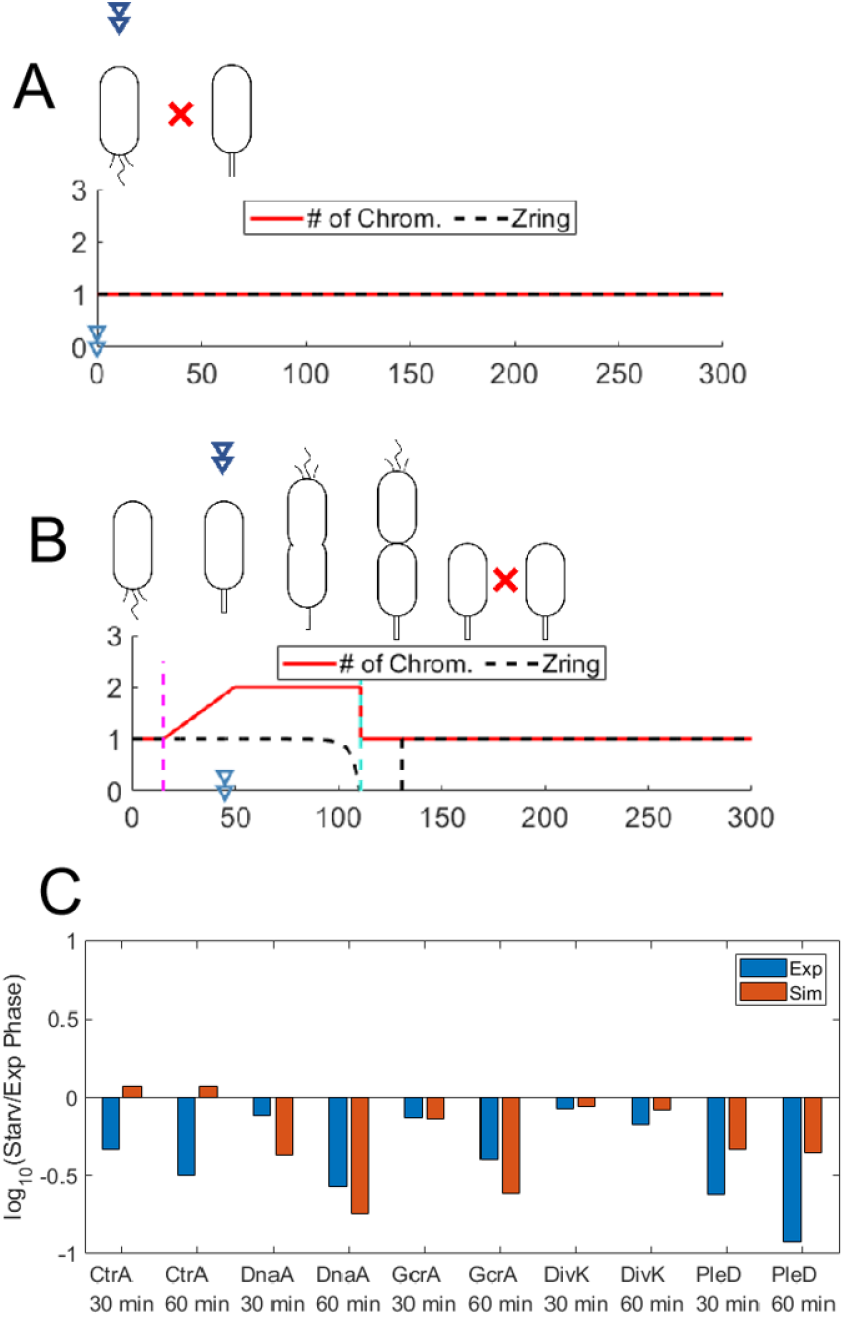
Cell cycle response of starvation Signal 1. (A) Swarmer cell simulation of Signal 1 response over time (min). Plot of chromosome count and Z-ring constriction. When the Zring variable is equal to 1, the Z-ring is fully open. It is closed when Zring is equal to 0. A double arrow indicates when the nutrient signal is introduced. A red “X” represents that swarmer cells arrest in the stage indicated (G1 phase for this figure). (B) Stalked cell simulation of Signal 1 response over time (min). (C) Protein expression levels 30 and 60 minutes after nutrient depletion (Signal 1) relative to nutrient-rich conditions.

**Figur 5:**
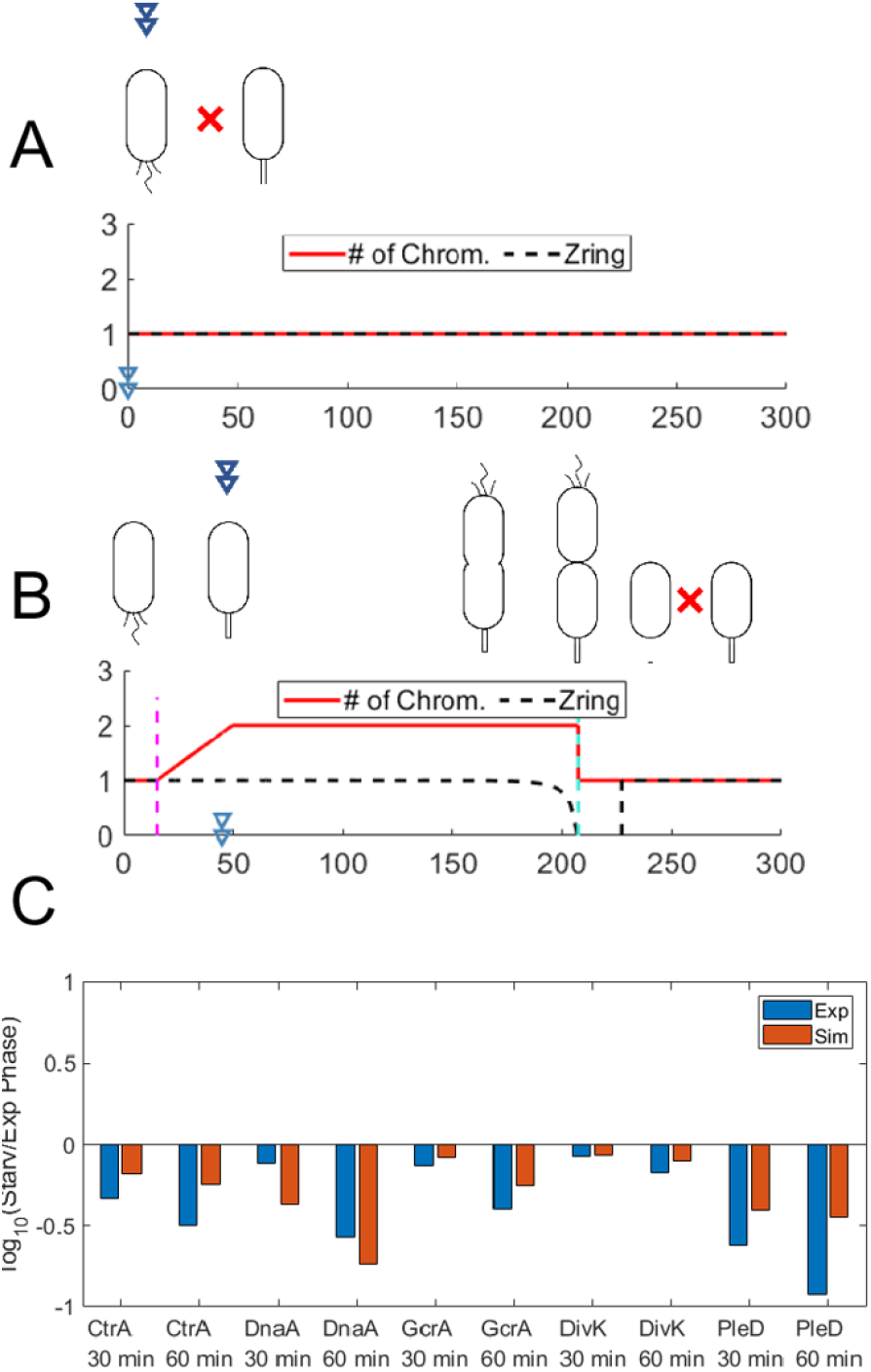
Starvation Signal 2 simulations fit experimental observations well with the reduction of CtrA expression. (A) Swarmer cell simulation of Signal 2 response over time (min). (B) Stalked cell simulation of Signal 2 response over time (min). (C) Protein expression levels 30 and 60 minutes after nutrient depletion relative to nutrient-rich conditions.

### 3.3 cdG response to starvation is essential but not sufficient to arrest the cell cycle

Here, we delete the cdG-dependent pathways from Signal 2, calling it Signal 3, and exclusively introduce reduced cdG levels and cell growth rate as Signal 4 to study the contribution of cdG in the response to starvation (Table 1). Without the cdG-dependent pathways, our simulated swarmer population exhibits secondary G1 arrest in 86.7% of our parameter sets, while only 13.3% of our parameter sets indicate immediate cell cycle arrest at G1 for swarmer cells (Table 1). This suggests that the cdG-involved second messenger network is significant for the immediate G1 arrest of swarmer cells under starvation. Additionally, we find that neither swarmer cells nor stalked cells arrest as experiments suggest in the simulation with Signal 4 via most parameter sets (Table 1), which demonstrates that cdG-dependent pathways are not sufficient to result in cell arrest in starved *Caulobacter* populations. Moreover, our simulation agrees with the experimental observation that cdG0 strain (deleting cdG) is not arrested.

## 4 Discussion

In this study, we investigate the molecular mechanism behind *Caulobacter* cell cycle arrest under nutrient starvation. Our model has good agreement with experimental observations in phenotypes of WT and mutant cells. Based on the rationality of this model, we further explore the contribution of known starvation-relevant pathways (Signal 1) and find these pathways are sufficient to capture most characteristics of starved cells except for two observed phenomena: 1) the delayed cytokinesis of starved stalked cells and 2) the reduced expression of CtrA under starvation. Therefore, we enforce the reduction of CtrA level in Signal 2, which solves the above two issues. This suggests that nutrient signals may influence CtrA via additional mechanisms to explain reduced CtrA expression under starvation.

Xu et al. [22] demonstrated that cdG levels are likely depleted due to the accumulation of (p)ppGpp, thus providing a link between (p)ppGpp regulation and CtrA proteolysis [22]. cdG is also responsible for the accumulation of SpmX at the G1-S transition via TacA and ShkA [9]. Modeling the ShkA-TacA-SpmX pathway reinforces the relationship between cdG and phosphorylation state of CtrA. We expand on this work here and find that the sole cdG response is not sufficient for cell cycle arrest (Table 1, Signal 4). However, removing the impaired cdG from the starvation signal failed to immediately and robustly arrest swarmer cells in G1 arrest (Table 1, Signal 3). Thus, while the proposed signal from Xu et al. [22] is not sufficient on its own to explain starvation response behaviors, it is an essential component of the starvation response mechanism.

Altogether, CtrA*∼*P remains high at the initial starvation stage because cdG is reduced, which is essential for the G1 arrest of starved cells. CtrA*∼*P level then decreases to delay the cytokinesis in starved stalked population, which is caused by an unclear control mechanism. Our results indicate that numerous aspects of the cell cycle machinery are targets of starvation signals. The phosphorylation of CtrA together with DnaA determines the G1 arrest, while CtrA also influences the timing of cytokinesis.

## References

[1] N. Ausmees and C. Jacobs-Wagner. Spatial and temporal control of differentiation and cell cycle progression in caulobacter crescentus. Annual Reviews in Microbiology, 57(1):225–247, 2003.

[2] E. G. Biondi, J. M. Skerker, M. Arif, M. S. Prasol, B. S. Perchuk, and M. T. Laub. A phospho-relay system controls stalk biogenesis during cell cycle progression in caulobacter crescentus. Molecular microbiology, 59(2):386–401, 2006.

[3] L. Britos, E. Abeliuk, T. Taverner, M. Lipton, H. McAdams, and L. Shapiro. Regulatory response to carbon starvation in caulobacter crescentus. PloS one, 6(4):e18179, 2011.

[4] L. C. B. Cavagnaro. Regulatory Response to Environmental Challenge in Caulobacter crescentus. Stanford University, 2011.

[5] B. N. Dubey, E. Agustoni, R. Böhm, A. Kaczmarczyk, F. Mangia, C. von Arx, U. Jenal, S. Hiller, I. Plaza-Menacho, and T. Schirmer. Hybrid histidine kinase activation by cyclic di-gmp–mediated domain liberation. Proceedings of the National Academy of Sciences, 117(2):1000–1008, 2020.

[6] B. Gorbatyuk and G. T. Marczynski. Regulated degradation of chromosome replication proteins dnaa and ctra in caulobacter crescentus. Molecular microbiology, 55(4):1233–1245, 2005.

[7] U. Jenal. The role of proteolysis in the caulobacter crescentus cell cycle and development. Research in microbiology, 160(9):687–695, 2009.

[8] K. K. Joshi, M. Bergé, S. K. Radhakrishnan, P. H. Viollier, and P. Chien. An adaptor hierarchy regulates proteolysis during a bacterial cell cycle. Cell, 163(2):419–431, 2015.

[9] A. Kaczmarczyk, A. M. Hempel, C. von Arx, R. Böhm, B. N. Dubey, J. Nesper, T. Schirmer, S. Hiller, and U. Jenal. Precise timing of transcription by c-di-gmp coordinates cell cycle and morphogenesis in caulobacter. Nature communications, 11(1):1–16, 2020.

[10] K. Lasker, T. H. Mann, and L. Shapiro. An intracellular compass spatially coordinates cell cycle modules in caulobacter crescentus. Current opinion in microbiology, 33:131–139, 2016.

[11] K. Lasker, J. M. Schrader, Y. Men, T. Marshik, D. L. Dill, H. H. McAdams, and L. Shapiro. Caulobrowser: A systems biology resource for caulobacter crescentus. Nucleic Acids Research, 44(D1):D640–D645, 2016.

[12] J. A. Lesley and L. Shapiro. Spot regulates dnaa stability and initiation of dna replication in carbon-starved caulobacter crescentus. Journal of bacteriology, 190(20):6867–6880, 2008.

[13] G. Li, C. S. Smith, Y. V. Brun, and J. X. Tang. The elastic properties of the caulobacter crescentus adhesive holdfast are dependent on oligomers of n-acetylglucosamine. Journal of bacteriology, 187(1):257–265, 2005.

[14] S. Li, P. Brazhnik, B. Sobral, and J. J. Tyson. A quantitative study of the division cycle of caulobacter crescentus stalked cells. PLoS computational biology, 4(1):e9, 2008.

[15] S. Li, P. Brazhnik, B. Sobral, and J. J. Tyson. Temporal controls of the asymmetric cell division cycle in caulobacter crescentus. PLoS computational biology, 5(8):e1000463, 2009.

[16] J. S. Poindexter. The caulobacters: ubiquitous unusual bacteria. Microbiological reviews, 45(1):123–179, 1981.

[17] S. Ronneau, K. Petit, X. De Bolle, and R. Hallez. Phosphotransferase-dependent accumulation of (p) ppgpp in response to glutamine deprivation in caulobacter crescentus. Nature communications, 7(1):1–12, 2016.

[18] S. C. Smith, K. K. Joshi, J. J. Zik, K. Trinh, A. Kamajaya, P. Chien, and K. R. Ryan. Cell cycle-dependent adaptor complex for clpxpmediated proteolysis directly integrates phosphorylation and second messenger signals. Proceedings of the National Academy of Sciences, 111(39):14229–14234, 2014.

[19] K. Subramanian, M. R. Paul, and J. J. Tyson. Potential role of a bistable histidine kinase switch in the asymmetric division cycle of caulobacter crescentus. PLoS computational biology, 9(9):e1003221, 2013.

[20] K. Subramanian, M. R. Paul, and J. J. Tyson. Dynamical localization of divl and plec in the asymmetric division cycle of caulobacter crescentus: a theoretical investigation of alternative models. PLoS computational biology, 11(7):e1004348, 2015.

[21] B. R. Weston, J. J. Tyson, and Y. Cao. Computational modeling of unphosphorylated ctra: Cori binding in the caulobacter cell cycle. Iscience, 24(12):103413, 2021.

[22] C. Xu, B. R. Weston, J. J. Tyson, and Y. Cao. Cell cycle control and environmental response by second messengers in caulobacter crescentus. BMC bioinformatics, 21(14):1–19, 2020.

