## Supplementary File 1 for "Modeling the Cell Cycle Response to Carbon and Nitrogen deprivation in *Caulobacter* Populations"

### Supplementary File 1: Equations of Modeling

Here, we provide the full set of equations for this study and all modifications compared with Weston et al.'s model are highlighted as red font.

Table 1: Equations.

(1) $$\frac{d[\mathrm{CtrA_U}]}{dt} = RpoD_{ctrA}\cdot k_{\mathrm{s,CtrA1}}\cdot(1 - m_{\mathrm{CtrA}}\cdot(2M_{\mathrm{ctrA}}-1))\cdot\frac{\epsilon_{CtrA-GcrA}\cdot J_{\mathrm{a,CtrAGcrA}}+[\mathrm{GcrA}]}{J_{\mathrm{a,CtrAGcrA}}+[\mathrm{GcrA}]}\cdot\left(1-\frac{[\mathrm{CtrA{\sim}P}]}{J_{\mathrm{i,CtrACtrA}}+[\mathrm{CtrA_U}]+[\mathrm{CtrA{\sim}P}]}\right) + k_{\mathrm{s,CtrA2}}\cdot\frac{\epsilon_{\mathrm{CtrACtrA}}\cdot J^2_{\mathrm{a,CtrACtrA{\sim}P}}+[\mathrm{CtrA{\sim}P}]^2}{J^2_{\mathrm{a,CtrACtrA{\sim}P}}+[\mathrm{CtrA_U}]^2+[\mathrm{CtrA{\sim}P}]^2}\cdot\frac{J^2_{\mathrm{i,CtrASciP}}}{J^2_{\mathrm{i,CtrASciP}}+[\mathrm{SciP}]^2} - \left(\mu + k_{\mathrm{d,CtrA1}} + k_{\mathrm{d,CtrA2}}\cdot\frac{[\mathrm{ClpXP}]_{\mathrm{complex}}}{J_{\mathrm{d,CtrA}}+[\mathrm{CtrA{\sim}P}]+[\mathrm{CtrA_U}]}\right)\cdot[\mathrm{CtrA_U}] + k_{\mathrm{dephos,CtrA}}\cdot[\mathrm{CckA_P}]\cdot[\mathrm{CtrA\sim P}] - k_{\mathrm{phos,CtrA}}\cdot[\mathrm{CtrA_U}]\cdot[\mathrm{CckA_K}]$$

(2) $$\frac{d[\mathrm{CtrA{\sim}P}]}{dt} = -\left(\mu + k_{\mathrm{d,CtrA1}} + k_{\mathrm{d,CtrA2}}\cdot\frac{[\mathrm{ClpXP}]_{\mathrm{complex}}}{J_{\mathrm{d,CtrA}}+[\mathrm{CtrA{\sim}P}]+[\mathrm{CtrA_U}]}\right)\cdot[\mathrm{CtrA\sim P}] - k_{\mathrm{dephos,CtrA}}\cdot[\mathrm{CckA_P}]\cdot[\mathrm{CtrA\sim P}] + k_{\mathrm{phos,CtrA}}\cdot[\mathrm{CtrA_U}]\cdot[\mathrm{CckA_K}]$$

(3) $$\frac{d[\mathrm{DnaA}]_{\mathrm{T}}}{dt} = k_{\mathrm{s,DnaA}}\cdot\frac{J_{\mathrm{i,DnaAGcrA}}}{J_{\mathrm{i,DnaAGcrA}}+[\mathrm{GcrA}]}\cdot(1 - 2\cdot m_{\mathrm{DnaA}}\cdot(1-M_{\mathrm{dnaA}})) - (\mu + k_{\mathrm{d,DnaA}})\cdot[\mathrm{DnaA}]_{\mathrm{t}}$$

(4) $$\frac{d[\mathrm{DnaA{\sim}ATP}]}{dt} = k_{\mathrm{s,DnaA}}\cdot\frac{J_{\mathrm{i,DnaAGcrA}}}{J_{\mathrm{i,DnaAGcrA}}+[\mathrm{GcrA}]}\cdot(1 - 2\cdot m_{\mathrm{DnaA}}\cdot(1-M_{\mathrm{dnaA}})) - (\mu + k_{\mathrm{d,DnaA}} + \mathrm{RepSwitch})\cdot[\mathrm{DnaA\sim ATP}]$$

(5) $$\frac{d[\mathrm{GcrA}]}{dt} = k_{\mathrm{s,GcrA}}\cdot\frac{\epsilon_{\mathrm{GcrADnaA}}\cdot J_{\mathrm{a,GcrADnaA}}+([\mathrm{DnaA}]_{\mathrm{T}}-[\mathrm{DnaA{\sim}ATP}])}{J_{\mathrm{a,GcrADnaA}}+([\mathrm{DnaA}]_{\mathrm{T}}-[\mathrm{DnaA{\sim}ATP}])}\cdot\frac{J^2_{\mathrm{i,GcrACtrA}}}{J^2_{\mathrm{i,GcrACtrA}}+[\mathrm{CtrA{\sim}P}]^2} - (\mu + k_{\mathrm{d,GcrA}})\cdot[\mathrm{GcrA}]$$

(6) $$\frac{d[\mathrm{SciP}]}{dt} = k_{\mathrm{s,SciP}}\cdot\frac{[\mathrm{CtrA{\sim}P}]^2}{[\mathrm{CtrA{\sim}P}]^2+J^2_{\mathrm{a,SciPCtrA}}}\cdot\frac{J^2_{\mathrm{i,SciPSciP}}}{J^2_{\mathrm{i,SciPSciP}}+[\mathrm{SciP}]^2} - (\mu + k_{\mathrm{d,SciP}})\cdot[\mathrm{SciP}]$$

(7) $$\frac{d[\mathrm{DivK}]}{dt} = RpoD_{divK}\cdot k_{\mathrm{s,DivK1}} + k_{\mathrm{s,DivK2}}\cdot\frac{[\mathrm{CtrA{\sim}P}]^2}{J^2_{\mathrm{a,DivKCtrA}}+[\mathrm{CtrA{\sim}P}]^2} - (\mu + k_{\mathrm{d,DivK}})\cdot[\mathrm{DivK}] - (k_{\mathrm{phos,DivK1}}\cdot[\mathrm{DivJ}]_{\mathrm{A}} + k_{\mathrm{phos,DivK2}}\cdot[\mathrm{PleC}]_{\mathrm{K}} + k_{\mathrm{phos,DivK3}}\cdot MysK)\cdot[\mathrm{DivK}] + (k_{\mathrm{dephos,DivK1}}\cdot[\mathrm{PleC}] + k_{\mathrm{dephos,DivK2}}\cdot[\mathrm{CckN}])\cdot[\mathrm{DivK\sim P}] - k_{\mathrm{b,DivJDivK}}\cdot[\mathrm{DivJ}]\cdot[\mathrm{DivK}] + (k_{\mathrm{ub,DivJDivK}} + k_{\mathrm{d,DivJ}})\cdot[\mathrm{DivJ:DivK}] - k_{\mathrm{b,DivLDivK}}\cdot[\mathrm{DivL}]\cdot[\mathrm{DivK}] + (k_{\mathrm{ub,DivLDivK}} + k_{\mathrm{d,DivL}})\cdot[\mathrm{DivL:DivK}]$$

(8) $$\frac{d[\mathrm{DivK{\sim}P}]}{dt} = -(\mu + k_{\mathrm{d,DivK}})\cdot[\mathrm{DivK}] - (k_{\mathrm{dephos,DivK1}}\cdot[\mathrm{PleC}] + k_{\mathrm{dephos,DivK2}}\cdot[\mathrm{CckN}])\cdot[\mathrm{DivK\sim P}] + (k_{\mathrm{phos,DivK1}}\cdot[\mathrm{DivJ}]_{\mathrm{A}} + k_{\mathrm{phos,DivK2}}\cdot[\mathrm{PleC}]_{\mathrm{K}} + k_{\mathrm{phos,DivK3}}\cdot MysK)\cdot[\mathrm{DivK}] + 2\cdot((k_{\mathrm{ub,PleCDivKP}} + k_{\mathrm{d,PleC}} + k_{\mathrm{d,DivK}})\cdot[\mathrm{PleC:DivK\sim P_2}] - k_{\mathrm{b,PleCDivKP}}\cdot[\mathrm{PleC}]\cdot[\mathrm{DivK\sim P}]^2) - k_{\mathrm{b,DivJDivKP}}\cdot[\mathrm{DivJ}]\cdot[\mathrm{DivK\sim P}] + (k_{\mathrm{ub,DivJDivKP}} + k_{\mathrm{d,DivJ}})\cdot[\mathrm{DivJ:DivK\sim P}] - k_{\mathrm{b,DivLDivKP}}\cdot[\mathrm{DivL}]\cdot[\mathrm{DivK\sim P}] + (k_{\mathrm{ub,DivLDivKP}} + k_{\mathrm{d,DivL}})\cdot[\mathrm{DivL:DivK\sim P}]$$

(9) $$\frac{d[\mathrm{CckN}]}{dt} = k_{\mathrm{s,CckN}}\cdot\frac{[\mathrm{CtrA{\sim}P}]^4}{[\mathrm{CtrA{\sim}P}]^4+J^4_{\mathrm{a,CckNCtrA}}} - (\mu + k_{\mathrm{d,CckN1}})\cdot[\mathrm{CckN}] - k_{\mathrm{d,CckN2}}\cdot[\mathrm{PopA:cdG_2}]\cdot\frac{[\mathrm{CckN}]}{[\mathrm{CckN}]+J_{\mathrm{d,CckN}}}$$

(10) $$\frac{[\mathrm{DivJ}]}{dt} = k_{\mathrm{s,DivJ}} - (\mu + k_{\mathrm{d,DivJ}})\cdot[\mathrm{DivJ}] - k_{\mathrm{b,DivJDivKP}}\cdot[\mathrm{DivJ}]\cdot[\mathrm{DivK\sim P}] + (k_{\mathrm{ub,DivJDivKP}} + k_{\mathrm{d,DivK}})\cdot[\mathrm{DivJ:DivK\sim P}] - k_{\mathrm{b,DivJDivK}}\cdot[\mathrm{DivJ}]\cdot[\mathrm{DivK}] + (k_{\mathrm{ub,DivJDivK}} + k_{\mathrm{d,DivK}})\cdot[\mathrm{DivJ:DivK}]$$

(11) $$\frac{d[\mathrm{DivJ{:}DivK}]}{dt} = k_{\mathrm{b,DivJDivK}}\cdot[\mathrm{DivJ}]\cdot[\mathrm{DivK}] - (k_{\mathrm{ub,DivJDivK}} + k_{\mathrm{d,DivK}} + +k_{\mathrm{d,DivJ}} + \mu)\cdot[\mathrm{DivJ:DivK}]$$

(11) $$\frac{d[\mathrm{DivJ{:}DivK{\sim}P}]}{dt} = k_{\mathrm{b,DivJDivK{\sim}P}}\cdot[\mathrm{DivJ}]\cdot[\mathrm{DivK\sim P}] - (k_{\mathrm{ub,DivJDivK{\sim}P}} + k_{\mathrm{d,DivK}} + +k_{\mathrm{d,DivJ}} + \mu)\cdot[\mathrm{DivJ:DivK\sim P}]$$

(12) $$[\mathrm{DivJ}]_{\mathrm{A}} = ([\mathrm{DivJ:DivK\sim P}] + [\mathrm{DivJ:DivK}])\cdot\left((1-\epsilon_{\mathrm{DivJDivK}})\cdot\left(\frac{\min([\mathrm{SpmX}],[\mathrm{DivJ}]_{\mathrm{T}})}{[\mathrm{DivJ}]_{\mathrm{T}}}\right) + \epsilon_{\mathrm{DivJDivK}}\right) + \epsilon_{\mathrm{DivJSpmX}}\cdot[\mathrm{DivJ}]\cdot\frac{\min([\mathrm{SpmX}],[\mathrm{DivJ}]_{\mathrm{T}})}{[\mathrm{DivJ}]_{\mathrm{T}}}$$

(13) $$\frac{[\mathrm{DivL}]}{dt} = k_{\mathrm{s,DivL}} - (\mu + k_{\mathrm{d,DivL}})\cdot[\mathrm{DivL}] - k_{\mathrm{b,DivLDivK{\sim}P}}\cdot[\mathrm{DivL}]\cdot[\mathrm{DivK\sim P}] + (k_{\mathrm{ub,DivLDivK{\sim}P}} + k_{\mathrm{d,DivK}})\cdot[\mathrm{DivL:DivK\sim P}]$$

(14) $$\frac{d[\mathrm{DivL{:}DivK{\sim}P}]}{dt} = k_{\mathrm{b,DivLDivK{\sim}P}}\cdot[\mathrm{DivL}]\cdot[\mathrm{DivK\sim P}] - (k_{\mathrm{ub,DivLDivK{\sim}P}} + k_{\mathrm{d,DivK}} + k_{\mathrm{d,DivL}} + \mu)\cdot[\mathrm{DivL:DivK\sim P}]$$

*Continued on next page*

Table A.1. – *Continued from previous page*

$$(15)\ \frac{[\mathrm{CckA}]_\mathrm{T}}{dt} = k_{\mathrm{s,CckA}} - (\mu + k_{\mathrm{d,CckA}}) \cdot [\mathrm{CckA}]_\mathrm{T}$$

$$(16)\ \frac{d[\mathrm{CckA:cdG}]}{dt} = k_{\mathrm{b,CckAcdG}} \cdot [\mathrm{cdG}] \cdot ([\mathrm{CckA}]_\mathrm{T} - [\mathrm{CckA:cdG}]) - (k_{\mathrm{ub,CckAcdG}} + k_{\mathrm{d,CckA}} + \mu) \cdot [\mathrm{CckA:cdG}]$$

$$(17)\ [\mathrm{CckA:DivL}]_\mathrm{T} = \frac{[\mathrm{CckA}]_\mathrm{T}+[\mathrm{DivL}]_\mathrm{T}+\frac{1}{K_{\mathrm{CckADivL}}} - \sqrt{([\mathrm{CckA}]_\mathrm{T}+[\mathrm{DivL}]_\mathrm{T}+\frac{1}{K_{\mathrm{CckADivL}}})^2 - 4\cdot[\mathrm{CckA}]_\mathrm{T}\cdot[\mathrm{DivL}]_\mathrm{T}}}{2}$$

$$(18)\ [\mathrm{CckA}]_\mathrm{P} = \frac{[\mathrm{CckA:DivL}]_\mathrm{T}+[\mathrm{DivL:DivK{\sim}P}]}{[\mathrm{DivL}]_\mathrm{T}} + [\mathrm{CckA:cdG}] - \frac{[\mathrm{CckA:cdG}]\cdot\frac{[\mathrm{CckA:DivL}]_\mathrm{T}\cdot[\mathrm{DivL:DivK{\sim}P}]}{[\mathrm{DivL}]_\mathrm{T}}}{[\mathrm{CckA}]_\mathrm{T}}$$

$$(19)\ [\mathrm{CckA}]_\mathrm{K} = [\mathrm{CckA}]_\mathrm{T} - [\mathrm{CckA}]_\mathrm{P}$$

$$(20)\ \frac{d[\mathrm{PleC}]}{dt} = k_{\mathrm{s,PleC}} \cdot (2 - 2\cdot M_{\mathrm{PleC}}) - (\mu + k_{\mathrm{d,PleC}}) \cdot [\mathrm{PleC}] - k_{\mathrm{b,PleCDivKP}} \cdot [\mathrm{PleC}] \cdot [\mathrm{DivK}\sim\mathrm{P}]^2 + (k_{\mathrm{ub,PleCDivKP}} + 2\cdot k_{\mathrm{d,DivK}})$$
$$\cdot[\mathrm{PleC:DivK}\sim\mathrm{P}_2]$$

$$(21)\ \frac{d[\mathrm{PleC:DivK{\sim}P_2}]}{dt} = k_{\mathrm{b,PleCDivKP}} \cdot [\mathrm{PleC}] \cdot [\mathrm{DivK}\sim\mathrm{P}]^2 - (k_{\mathrm{ub,PleCDivKP}} + 2\cdot k_{\mathrm{d,DivK}} + k_{\mathrm{d,PleC}} + \mu) \cdot [\mathrm{PleC:DivK}\sim\mathrm{P}_2]$$

$$(22)\ \frac{d[\mathrm{PleC}]_{\mathrm{pole}}}{dt} = k_{\mathrm{PleCbinding}} \cdot ([\mathrm{PleC}]_\mathrm{T} - [\mathrm{PleC}]_{\mathrm{pole}}) \cdot \frac{[\mathrm{PodJ}]}{[\mathrm{PodJ}]\cdot V + J_{\mathrm{PleCPodJ}}} - (k_{\mathrm{PleCbinding}} + k_{\mathrm{d,PleC}} + \mu) \cdot [\mathrm{PleC}]_{\mathrm{pole}}$$

$$(23)\ \frac{d[\mathrm{PodJ}]}{dt} = k_{\mathrm{s,PodJ}} \cdot \frac{[\mathrm{GcrA}]}{[\mathrm{GcrA}]+J_{\mathrm{a,PodJGcrA}}} \cdot \frac{J_{\mathrm{i,PodJDnaA}}}{J_{\mathrm{i,PodJDnaA}}+[\mathrm{PodJ}]} \cdot (2 - 2\cdot M_{\mathrm{PodJ}}) - (\mu + k_{\mathrm{d,PodJ1}} + k_{\mathrm{d,PodJ2}} \cdot [\mathrm{PerP}]) \cdot [\mathrm{PodJ}]$$

$$(24)\ \frac{d[\mathrm{PerP}]}{dt} = k_{\mathrm{s,PerP}} \cdot \frac{[\mathrm{CtrA}]^2}{[\mathrm{CtrA}]^2+J^2_{\mathrm{a,PerPCtrA}}} \cdot (2 - 2\cdot M_{\mathrm{PerP}}) - (\mu + k_{\mathrm{d,PerP}}) \cdot [\mathrm{PerP}]$$

$$(25)\ \frac{d[\mathrm{Ini}]}{dt} = (1 - 2\cdot m_{\mathrm{ini}} \cdot (1 - M_{\mathrm{Ini}})) \cdot (1 - [\mathrm{DNA:CtrA}\sim\mathrm{P}_2])^5 \cdot \left(\frac{[\mathrm{DNA{\sim}ATP}]}{[\mathrm{DNA{\sim}ATP}]+J_{\mathrm{a,IniDnaA}}}\right)^2 - k_{\mathrm{d,Ini}} \cdot [\mathrm{Ini}]$$

$$(26)\ [\mathrm{DNA}]_\mathrm{F} = \frac{K_{\mathrm{d1}}}{K_{\mathrm{d1}} + 2\cdot\mathrm{CtrA{\sim}P} + (1-\sigma_{\mathrm{CtrAU}:Cori})\cdot(2\cdot[\mathrm{CtrA_U}] + \frac{[\mathrm{CtrA_U}]^2}{K_{\mathrm{d2}}} + \frac{[\mathrm{CtrA_U}]+[\mathrm{CtrA{\sim}P}]}{K_{\mathrm{d2}}})}$$

$$(27)\ [\mathrm{DNA:CtrA}\sim\mathrm{P}_2] = \frac{[\mathrm{CtrA{\sim}P}]^2\cdot[\mathrm{DNA}]_\mathrm{F}}{K_{\mathrm{d1}}\cdot K_{\mathrm{d3}}}$$

$$(28)\ \frac{d[\mathrm{Elong}]}{dt} = k_{\mathrm{elong}} \cdot RepSwitch$$

$$(29)\ \frac{d[\mathrm{CcrM}]}{dt} = k_{\mathrm{s,CcrM}} \cdot \frac{[\mathrm{CtrA}]^2}{[\mathrm{CtrA}]^2+J^2_{\mathrm{a,CcrMCtrA}}} \cdot \frac{J_{\mathrm{i,CcrMDnaA}}}{J_{\mathrm{i,CcrMDnaA}}+[\mathrm{DnaA}]} \cdot (2 - 2\cdot M_{\mathrm{CcrM}}) - (\mu + k_{\mathrm{d,CcrM}}) \cdot [\mathrm{CcrM}]$$

$$(30)\ \frac{d[\mathrm{TacA}]}{dt} = k_{\mathrm{s,TacA}} \cdot \frac{[\mathrm{CtrA{\sim}P}]}{[\mathrm{CtrA{\sim}P}]+J_{\mathrm{a,TacACtrA}}} - (k_{\mathrm{d,TacA1}} + \mu) \cdot [\mathrm{TacA}] - k_{\mathrm{d,TacA2}} \cdot [\mathrm{RcdA:CpdR}] \cdot \frac{[\mathrm{TacA}]}{[\mathrm{TacA}]_\mathrm{T}+J_{\mathrm{d,TacA}}}$$
$$-k_{\mathrm{phos,TacA}} \cdot [\mathrm{ShkA:cdG}] \cdot [\mathrm{TacA}] + k_{\mathrm{dephos,TacA}} \cdot [\mathrm{ShkA}] \cdot [\mathrm{TacA}\sim\mathrm{P}]$$

$$(31)\ \frac{d[\mathrm{TacA{\sim}P}]}{dt} = -k_{\mathrm{d,TacA2}} \cdot [\mathrm{RcdA:CpdR}] \cdot \frac{[\mathrm{TacA{\sim}P}]}{[\mathrm{TacA}]_\mathrm{T}+J_{\mathrm{d,TacA}}} + k_{\mathrm{phos,TacA}} \cdot [\mathrm{ShkA:cdG}] \cdot [\mathrm{TacA}]$$
$$-(\mu + k_{\mathrm{d,TacA1}} + k_{\mathrm{dephos,TacA}} \cdot [\mathrm{ShkA}]) \cdot [\mathrm{TacA}\sim\mathrm{P}]$$

$$(32)\ \frac{d[\mathrm{SpmX}]}{dt} = k_{\mathrm{s,SpmX}} \cdot \frac{[\mathrm{TacA{\sim}P}]}{[\mathrm{TacA{\sim}P}]+J_{\mathrm{a,SpmXTacA}}} - (k_{\mathrm{d,SpmX}} + \mu) \cdot [\mathrm{SpmX}]$$

$$(33)\ \frac{d[\mathrm{ShkA}]}{dt} = k_{\mathrm{s,ShkA}} \cdot \frac{[\mathrm{CtrA{\sim}P}]}{[\mathrm{CtrA{\sim}P}]+J_{\mathrm{a,ShkACtrA}}} - (k_{\mathrm{d,ShkA1}} + \mu) \cdot [\mathrm{ShkA}] - k_{\mathrm{d,ShkA2}} \cdot [\mathrm{ClpXP}]_{\mathrm{Complex}} \cdot \frac{[\mathrm{ShkA}]}{[\mathrm{ShkA}]_\mathrm{T}+J_{\mathrm{d,ShkA}}}$$
$$-k_{\mathrm{b,ShkAcdG}} \cdot [\mathrm{cdG}] \cdot [\mathrm{ShkA}] + k_{\mathrm{ub,ShkAcdG}} \cdot [\mathrm{ShkA:cdG}]$$

$$(34)\ \frac{d[\mathrm{ShkA:cdG}]}{dt} = -(\mu + k_{\mathrm{d,ShkA1}}) \cdot [\mathrm{ShkA:cdG}] - k_{\mathrm{d,ShkA2}} \cdot [\mathrm{ClpXP}]_{\mathrm{Complex}} \cdot \frac{[\mathrm{ShkA:cdG}]}{[\mathrm{ShkA}]_\mathrm{T}+J_{\mathrm{d,ShkA}}}$$
$$+k_{\mathrm{b,ShkAcdG}} \cdot [\mathrm{cdG}] \cdot [\mathrm{ShkA}] - k_{\mathrm{ub,ShkAcdG}} \cdot [\mathrm{ShkA:cdG}]$$

$$(35)\ \frac{d\mathrm{Zproteins}}{dt} = k_{\mathrm{s,Zp}} \cdot \frac{[\mathrm{CtrA}]^2}{[\mathrm{CtrA}]^2+J^2_{\mathrm{a,ZpCtrA}}} - (\mu + k_{\mathrm{d,Zp1}} + k_{\mathrm{d,Zp2}} \cdot [\mathrm{ClpAP}]) \cdot [\mathrm{Zproteins}]$$

$$(36)\ \frac{d[\mathrm{Zring}]}{dt} = -k_{\mathrm{Zconstrict}} \cdot MipZswitch \cdot \frac{[\mathrm{Zproteins}]^5}{(J_{\mathrm{Zring}}+\theta_\mathrm{Z}\cdot[\mathrm{Zring}])^5+[\mathrm{Zproteins}]^5}$$

$$(37)\ \frac{dV}{dt} = \mu V$$

$$(38)\ \mu = T_{-1} \cdot ln\frac{V_{\mathrm{div}}}{V_{\mathrm{birth}}}$$

$$(39)\ \frac{d[\mathrm{CpdR}]}{dt} = k_{\mathrm{s,CpdR}} \cdot \frac{\epsilon_{\mathrm{CpdRDnaA}}\cdot J_{\mathrm{a,CpdRDnaA}}+[\mathrm{DnaA}]_\mathrm{T}}{J_{\mathrm{a,CpdRDnaA}}+[\mathrm{DnaA}]_\mathrm{T}} \cdot \frac{[\mathrm{CtrA}]^2}{[\mathrm{CtrA}]^2+J^2_{\mathrm{a,CpdRCtrA}}} \cdot \frac{J_{\mathrm{i,CpdRGcrA}}}{J_{\mathrm{i,CpdRGcrA}}+[\mathrm{GcrA}]} - (\mu + k_{\mathrm{d,CpdR}}) \cdot [\mathrm{CpdR}]$$
$$+k_{\mathrm{dephos,CpdR}} \cdot [\mathrm{CpdR}\sim\mathrm{P}] \cdot [\mathrm{CckA}]_\mathrm{P} - k_{\mathrm{phos,CpdR}} \cdot [\mathrm{CpdR}] \cdot [\mathrm{CckA}]_\mathrm{K}$$

$$(40)\ \frac{d[\mathrm{CpdR{\sim}P}]}{dt} = -k_{\mathrm{dephos,CpdR}} \cdot [\mathrm{CpdR}\sim\mathrm{P}] \cdot [\mathrm{CckA}]_\mathrm{P} + k_{\mathrm{phos,CpdR}} \cdot [\mathrm{CpdR}] \cdot [\mathrm{CckA}]_\mathrm{K}$$

$$(41)\ \frac{d[\mathrm{RcdA}]}{dt} = k_{\mathrm{s,RcdA}} \cdot \frac{[\mathrm{CtrA{\sim}P}]^2}{[\mathrm{CtrA{\sim}P}]^2+J^2_{\mathrm{a,RcdACtrA}}} - (\mu + k_{\mathrm{d,RcdA}}) \cdot [\mathrm{RcdA}]$$

$$(42)\ \frac{d[\mathrm{cdG}]}{dt} = (k_{\mathrm{s,cdG1}} \cdot [\mathrm{PleC}\sim\mathrm{P}] + k_{\mathrm{s,cdG2}} \cdot [\mathrm{DgcB}]_\mathrm{a}) \cdot \frac{[\mathrm{GTP}]^2}{[\mathrm{GTP}]^2+J^2_{\mathrm{s,cdG}}} - k_{\mathrm{d,cdG1}} \cdot ([\mathrm{PdeA}] + PDE) \cdot [\mathrm{cdG}] - \mu \cdot [\mathrm{cdG}]$$
$$+2\cdot(-k^+_{\mathrm{PopAcdG}} \cdot [\mathrm{PopA}] \cdot [\mathrm{cdG}]^2 + (k^-_{\mathrm{XcdG}} + K_{\mathrm{d,PopA}}) \cdot [\mathrm{PopA:cdG_2}]) + 2\cdot(-k^+_{\mathrm{PleDcdG}} \cdot [\mathrm{PleD}]_\mathrm{T} \cdot [\mathrm{cdG}]^2$$

*Continued on next page*

Table A.1. – *Continued from previous page*

$+(k^{-}_{\mathrm{XcdG}} + K_{\mathrm{d,PleD}}) \cdot [\mathrm{PleD:cdG_2}]_{\mathrm{T}}) - k^{+}_{\mathrm{CckAcdG}} \cdot ([\mathrm{CckA}]_{\mathrm{T}} - [\mathrm{CckA:cdG}]) \cdot [\mathrm{cdG}] + k^{-}_{\mathrm{CckAcdG}} \cdot [\mathrm{CckA:cdG}]$

$+2 \cdot (-k^{+}_{\mathrm{DgcBcdG}} \cdot (DgcB - [\mathrm{DgcB:cdG_2}])) \cdot [\mathrm{cdG}]^2 + k^{-}_{\mathrm{XcdG}} \cdot [\mathrm{DgcB:cdG_2}]$

(43) $\frac{d[\mathrm{PopA}]}{dt} = k_{\mathrm{s,PopA}} \cdot \frac{J_{\mathrm{i,PopAGcrA}}}{J_{\mathrm{i,PopAGcrA}}+[\mathrm{GcrA}]} - (\mu + k_{\mathrm{d,PopA}}) \cdot [\mathrm{PopA}] - k_{\mathrm{b,PopAcdG}} \cdot [\mathrm{PopA}] \cdot [\mathrm{cdG}]^2 + k_{\mathrm{ub,PopAcdG}} \cdot [\mathrm{PopA:cdG_2}]$

(44) $\frac{d[\mathrm{PopA:cdG_2}]}{dt} = k_{\mathrm{b,PopAcdG}} \cdot [\mathrm{PopA}] \cdot [\mathrm{cdG}]^2 - (\mu + k_{\mathrm{d,PopA}} + k_{\mathrm{ub,PopAcdG}}) \cdot [\mathrm{PopA:cdG_2}]$

(45) $[\mathrm{RcdA:CpdR}]_{\mathrm{T}} = \frac{[\mathrm{CpdR}]_{\mathrm{T}}+[\mathrm{RcdA}]+\frac{1}{K_{\mathrm{RcdACpdR}}} - \sqrt{([\mathrm{CpdR}]_{\mathrm{T}}+[\mathrm{RcdA}]+\frac{1}{K_{\mathrm{RcdACpdR}}})^2 - 4 \cdot [\mathrm{CpdR}]_{\mathrm{T}} \cdot [\mathrm{RcdA}]}}{2}$

(46) $[\mathrm{RcdA:CpdR}] = [\mathrm{RcdA:CpdR}]_{\mathrm{T}} \cdot \frac{[\mathrm{CpdR}]}{[\mathrm{CpdR}]_{\mathrm{T}}}$

(47) $[\mathrm{ClpXP}]_{\mathrm{Complex}} = \frac{[\mathrm{RcdA:CpdR}]}{[\mathrm{RcdA:CpdR}]+\frac{K_{\mathrm{ClpXPCpdR}}}{V}} \cdot [\mathrm{PopA:cdG_2}]$

(48) $\frac{d[\mathrm{PdeA}]}{dt} = k_{\mathrm{s,PdeA}} \cdot \frac{[\mathrm{CtrA{\sim}P}]}{[\mathrm{CtrA{\sim}P}]+J_{\mathrm{a,PdeACtrA}}} - (\mu + k_{\mathrm{d,PdeA1}}) \cdot [\mathrm{PdeA}] - k_{\mathrm{d,PdeA2}} \cdot \frac{[\mathrm{PdeA}]}{[\mathrm{PdeA}]+J_{\mathrm{d,PdeA}}}$

(49) $[\mathrm{DgcB}]_{\mathrm{a}} = \max(DgcB - \mathrm{PdeA}, 0) \cdot \frac{DgcB - [\mathrm{DgcB:cdG_2}]}{DgcB}$

(50) $\frac{[\mathrm{DgcB:cdG_2}]}{dt} = k_{\mathrm{b,DgcBcdG}} \cdot (DgcB - [\mathrm{DgcB:cdG_2}]) \cdot [\mathrm{cdG}]^2 - (\mu + k_{\mathrm{ub,XcdG}}) \cdot [\mathrm{DgcB:cdG_2}]$

(51) $\frac{d[\mathrm{PleD}]}{dt} = k_{\mathrm{s,PleD1}} \cdot \frac{[\mathrm{CtrA{\sim}P}]^2}{[\mathrm{CtrA{\sim}P}]^2+J^2_{\mathrm{a,PleDCtrA}}} + k_{\mathrm{s,PleD2}} \cdot RpoD_{pleD} - (\mu + k_{\mathrm{d,PleD}}) \cdot [\mathrm{PleD}] - k_{\mathrm{phos,PleD}} \cdot ([\mathrm{DivJ}]_{\mathrm{A}}$

$+MysK) \cdot [\mathrm{PleD}] + (\frac{1}{10} + \frac{9}{10} \cdot \frac{[\mathrm{PleC}]_{\mathrm{tot}} - [\mathrm{PleC}]_{\mathrm{pole}}}{[\mathrm{PleC}]_{\mathrm{tot}}}) \cdot k_{\mathrm{dephos,PleCPleD}} \cdot [\mathrm{PleC}] \cdot [\mathrm{PleD} \sim \mathrm{P}] + k_{\mathrm{dephos,CckNPleD}} \cdot [\mathrm{CckN}]$

$-k_{\mathrm{b,PleDcdG}} \cdot [\mathrm{PleD}] \cdot [\mathrm{cdG}]^2 + k_{\mathrm{ub,PleDcdG}} \cdot [\mathrm{PleD:cdG_2}]$

(52) $\frac{d[\mathrm{PleD{\sim}P}]}{dt} = -(\mu + k_{\mathrm{d,PleD}}) \cdot [\mathrm{PleD}] + k_{\mathrm{phos,PleD}} \cdot ([\mathrm{DivJ}]_{\mathrm{A}} + MysK) \cdot [\mathrm{PleD}] - (\frac{1}{10} + \frac{9}{10} \cdot \frac{[\mathrm{PleC}]_{\mathrm{tot}} - [\mathrm{PleC}]_{\mathrm{pole}}}{[\mathrm{PleC}]_{\mathrm{tot}}})$

$\cdot k_{\mathrm{dephos,PleCPleD}} \cdot [\mathrm{PleC}] \cdot [\mathrm{PleD} \sim \mathrm{P}] - k_{\mathrm{dephos,CckNPleD}} \cdot [\mathrm{CckN}] \cdot [\mathrm{PleD} \sim \mathrm{P}]$

$-k_{\mathrm{b,PleDcdG}} \cdot [\mathrm{PleD} \sim \mathrm{P}] \cdot [\mathrm{cdG}]^2 + k_{\mathrm{ub,PleDcdG}} \cdot [\mathrm{PleD} \sim \mathrm{P:cdG_2}]$

(53) $\frac{d[\mathrm{PleD:cdG_2}]}{dt} = -(\mu + k_{\mathrm{d,PleD}}) \cdot [\mathrm{PleD:cdG_2}] - k_{\mathrm{phos,PleD}} \cdot ([\mathrm{DivJ}]_{\mathrm{A}} + MysK) \cdot [\mathrm{PleD:cdG_2}] + (\frac{1}{10} + \frac{9}{10} \cdot \frac{[\mathrm{PleC}]_{\mathrm{tot}} - [\mathrm{PleC}]_{\mathrm{pole}}}{[\mathrm{PleC}]_{\mathrm{tot}}})$

$\cdot k_{\mathrm{dephos,PleCPleD}} \cdot [\mathrm{PleC}] \cdot [\mathrm{PleD} \sim \mathrm{P:cdG_2}] + k_{\mathrm{dephos,CckNPleD}} \cdot [\mathrm{CckN}] \cdot [\mathrm{PleD} \sim \mathrm{P:cdG_2}] + k_{\mathrm{b,PleDcdG}} \cdot [\mathrm{PleD}] \cdot [\mathrm{cdG}]^2$

$-k_{\mathrm{ub,PleDcdG}} \cdot [\mathrm{PleD:cdG_2}]$

(54) $\frac{d[\mathrm{PleD{\sim}P:cdG_2}]}{dt} = -(\mu + k_{\mathrm{d,PleD}}) \cdot [\mathrm{PleD} \sim \mathrm{P:cdG_2}] + k_{\mathrm{phos,PleD}} \cdot ([\mathrm{DivJ}]_{\mathrm{A}} + MysK) \cdot [\mathrm{PleD:cdG_2}]$

$-(\frac{1}{10} + \frac{9}{10} \cdot \frac{[\mathrm{PleC}]_{\mathrm{tot}} - [\mathrm{PleC}]_{\mathrm{pole}}}{[\mathrm{PleC}]_{\mathrm{tot}}}) \cdot k_{\mathrm{dephos,PleCPleD}} \cdot [\mathrm{PleC}] \cdot [\mathrm{PleD} \sim \mathrm{P:cdG_2}] - k_{\mathrm{dephos,CckNPleD}} \cdot [\mathrm{CckN}] \cdot [\mathrm{PleD} \sim \mathrm{P:cdG_2}]$

$-k_{\mathrm{ub,PleDcdG}} \cdot [\mathrm{PleD} \sim \mathrm{P:cdG_2}] + k_{\mathrm{b,PleDcdG}} \cdot [\mathrm{PleD} \sim \mathrm{P}] \cdot [\mathrm{cdG}]^2$
