## Supplementary File 2 for "Modeling the Cell Cycle Response to Carbon and Nitrogen deprivation in *Caulobacter* Populations"

### Supplementary File 2: Cost Functions of Mutant Strains

***ΔccrM***:

$$S_{\Delta ccrM,\mathrm{CT}} = \left(\frac{150}{N_{\mathrm{cycle}}}\right)^2, \text{ if cell cycle arrested,} \tag{1}$$

$$S^c_{\Delta ccrM,\mathrm{SW}} = \left(\frac{t_{\mathrm{div,c}} - 180}{15}\right)^2, \text{ if cell cycle not arrested,} \tag{2}$$

$$S^c_{\Delta ccrM,\mathrm{ST}} = 0, \quad \text{if cell cycle not arrested,} \tag{3}$$

***ΔgcrA***:

$$S_{\Delta gcrA,\mathrm{CT}} = \left(\frac{150}{N_{\mathrm{cycle}}}\right)^2, \text{ if cell cycle arrested,} \tag{4}$$

$$S^c_{\Delta gcrA,\mathrm{SW}} = \left(\frac{t_{\mathrm{div,c}} - 180}{8}\right)^2, \text{ if cell cycle not arrested,} \tag{5}$$

$$S^c_{\Delta gcrA,\mathrm{ST}} = 0, \quad \text{if cell cycle not arrested,} \tag{6}$$

***ΔpleD***:

$$S_{\Delta pleD,\mathrm{CT}} = \left(\frac{150}{N_{\mathrm{cycle}}}\right)^2, \text{ if cell cycle arrested,} \tag{7}$$

$$S^c_{\Delta pleD,\mathrm{SW}} = \left(\frac{t_{\mathrm{div,c}} - 145}{5}\right)^2 + \left(\frac{t_{\mathrm{rep,c}} - 20}{4}\right)^2 + w_{\mathrm{CtrA}} \cdot S_{\mathrm{CtrA}}, \text{ if cell cycle not arrested,} \tag{8}$$

$$S^c_{\Delta pleD,\mathrm{ST}} = \left(\frac{t_{\mathrm{div,c}} - 115}{5}\right)^2, \text{ if cell cycle not arrested,} \tag{9}$$

***divLA601L*** :

$$S^{c}_{divL\text{A601A,CT}} = \frac{5\times10^4}{t_{\text{div},c}}, \text{ if cell cycle not arrested,} \tag{10}$$

$$S^{c}_{divL\text{A601A,CT}} = 5\times(N_{cycles}-1), \text{ if cell cycle arrested,} \tag{11}$$

***ctrA*Δ3Ω**:

$$S_{ctrA\Delta3\Omega\text{,CT}} = \left(\frac{150}{N_{\text{cycle}}}\right)^2, \text{ if cell cycle arrested,} \tag{12}$$

$$S^{c}_{ctrA\Delta3\Omega\text{,SW}} = 2\cdot\left(\frac{t_{\text{div,c}}-145}{5}\right)^2+\left(\frac{t_{\text{rep,c}}-20}{4}\right)^2, \text{ if cell cycle not arrested,} \tag{13}$$

$$S^{c}_{ctrA\Delta3\Omega\text{,ST}} = 2\cdot\left(\frac{t_{\text{div,c}}-115}{5}\right)^2, \text{ if cell cycle not arrested,} \tag{14}$$

***ctrAD51E*** :

$$S_{ctrA\text{D51E,CT}} = \left(\frac{150}{N_{\text{cycle}}}\right)^2, \text{ if cell cycle arrested,} \tag{15}$$

$$S^{c}_{ctrA\text{D51E,SW}} = \left(\frac{t_{\text{div,c}}-145}{5}\right)^2+\left(\frac{t_{\text{rep,c}}-20}{4}\right)^2, \text{ if cell cycle not arrested,} \tag{16}$$

$$S^{c}_{ctrA\text{D51E,ST}} = \left(\frac{\text{t}_{\text{div,c}}-115}{5}\right)^2, \text{ if cell cycle not arrested,} \tag{17}$$

***PpleC*:: *Tn*&Δ*divJ***:

$$S_{\text{P}pleC\text{::Tn\&}\Delta DIVj\text{,CT}} = \left(\frac{1500}{N_{\text{cycle}}}\right)^2, \text{ if cell cycle arrested,} \tag{18}$$

$$S_{\text{P}pleC\text{::Tn\&}\Delta DIVj\text{,CT}} = 0, \text{ if cell cycle not arrested,} \tag{19}$$
